# Pat1 activates late steps in mRNA decay by multiple mechanisms

**DOI:** 10.1101/594168

**Authors:** Joseph H. Lobel, Ryan W. Tibble, John D. Gross

## Abstract

Pat1 is a hub for mRNA metabolism, acting in pre-mRNA splicing, translation repression and mRNA decay. A critical step in all 5’-3’ mRNA decay pathways is removal of the 5’ cap structure, which precedes and permits digestion of the RNA body by conserved exonucleases. During bulk 5’-3’ decay, the Pat1/Lsm1-7 complex engages mRNA at the 3’ end and promotes hydrolysis of the cap structure by Dcp1/Dcp2 at the 5’ end through an unknown mechanism. We reconstitute Pat1 with 5’ and 3’ decay factors and show how it activates multiple steps in late mRNA decay. First, we find that Pat1 stabilizes binding of the Lsm1-7 complex to RNA using two conserved short-linear interaction motifs. Secondly, Pat1 directly activates decapping by binding elements in the disordered C-terminal extension of Dcp2, alleviating autoinhibition and promoting substrate binding. Our results uncover the molecular mechanism of how separate domains of Pat1 coordinate the assembly and activation of a decapping mRNP that promotes 5’-3’ mRNA degradation.

## INTRODUCTION

Proper degradation of mRNA transcripts shapes the timing and quantity of gene expression to regulate a diverse array of cellular processes (1, 2). Cleavage of the 5’ m^7^G cap (decapping) removes the mRNA from the translating pool and exposes a free monophosphate which leads to rapid degradation by the 5’-3’ exonuclease Xrn1(2, 3). Decapping is a critical process for establishing 5’-3’ mRNA degradation and is found in bulk mRNA decay, clearance of maternal transcripts, quality control pathways, and miRNA mediated decay (4–7). The Dcp1/Dcp2 holoenzyme is a conserved NUDIX hydrolase that cleaves the 5’ m^7^G cap and is targeted to specific mRNAs by cofactors (1, 8–13). These cofactors regulate the activity of Dcp1/Dcp2 by either binding the catalytic core of the enzyme or helical leucine motifs (HLMs) in the disordered C-terminus of Dcp2, which contains additional *cis-*regulatory elements that inhibit decapping (14–17). Thus, a dense network of protein interactions has evolved to coordinate decapping of specific mRNA targets (18).

Bulk 5’-3’ mRNA degradation begins with trimming of the 3’ poly(A) tail, which may be followed by the addition of 1-3 uridines by terminal urdiyl transferases in higher order eukaryotes (19, 20). Deadenylation results in loss of translation initiation factors and permits the assembly of a decapping mRNP (21). Pat1 and Lsm1-7 work as a complex that assembles at the 3’ end of an mRNA to subsequently promote decapping by Dcp1/Dcp2 (21–26). Structural and biochemical studies have demonstrated that Pat1 and Lsm1-7 form a heterooctameric complex that engages transcripts containing oligo(A) tails that result from deadenylation (27–30). Pat1 and Lsm1-7 are functionally linked, as they bind the 3’ end of mRNA and deletion of either Pat1 or Lsm1 causes an increase in steady-state levels of numerous overlapping transcripts (12, 30). In addition, deletion of Pat1 or Lsm1 results in an accumulation of deadenylated, capped mRNA intermediates, suggesting a block in decapping (21, 22, 31). Finally, Pat1 forms a core component of P-bodies and can recruit interacting partners to these cytoplasmic foci, which may function in mRNA storage or decay (32–34). A critical and unresolved question is how Pat1 coordinates assembly of Lsm1-7 complex on the mRNA 3’ end with decapping at the 5’ end.

Pat1 is conserved from yeast to humans and contains three domains that interact with distinct decay factors. The disordered N-terminal region of Pat1 contains a conserved motif that interacts with the DEAD-box ATPase Dhh1, but is generally dispensable for cell growth and normal mRNA turnover (35, 36). The α-α superhelical C-terminal domain (PatC) interacts with Lsm1-7, which is required for bulk 5’-3’ mRNA degradation *in vivo* (27, 28, 37–39). Removal of the unstructured middle domain severely attenuates decapping and 5’-3’ degradation, though the mechanism remains poorly understood (36, 40). The conservation of the middle and C-terminal domains from yeast to humans suggests they are critical for decapping and proper mRNA decay, but how these domains interact with and activate mRNA decay factors remains unknown.

Here, we determine how Pat1 interacts with and activates both the Lsm1-7 complex and Dcp1/Dcp2 from the fission yeast *S.pombe (Sp)* using recombinant purified proteins, which recapitulate the regulation of decapping observed in budding yeast (16, 17). We show that Pat1 promotes RNA binding of the Lsm1-7 complex using two short linear motifs from its middle domain that make physical interactions with the Lsm1-7 ring and enhance its interaction with RNA. Pat1 activates decapping using its middle domain and structured C-terminus, which directly binds conserved helical leucine motifs in Dcp2, to alleviate autoinhibition and promote substrate binding. Our results reveal how Pat1 nucleates assembly of a decapping mRNP and uses distinct domains to activate decay factors at both the 3’ and 5’ ends of a transcript to promote mRNA degradation.

## RESULTS

### Pat1 makes bipartite interactions with Lsm1-7 to enhance RNA binding

Previous studies of Pat1 and Lsm1-7 purified from budding yeast indicate Pat1 is necessary to promote high affinity interactions with Lsm1-7 and RNA, and that the middle and C-terminal domains of Pat1 are sufficient to complement growth defects in strains where Pat1 is deleted (36, 37). To understand the domains of Pat1 involved in mRNA turnover in S. *pombe*, we constructed strains harboring different domain deletions of Pat1(Fig. 1A). Both full length Pat1 and PatMC (residues 296-754) were sufficient to compliment the growth defect in the ∆Pat1 strain. Neither the middle nor C-terminal domain alone, however, could rescue growth, which was not due to a protein expression defect (Fig. 1B,C). This suggests that, like budding yeast, both the middle and C-terminal domains of Pat1 are required for normal mRNA turnover in S.*pombe*.

**Figure 1:**
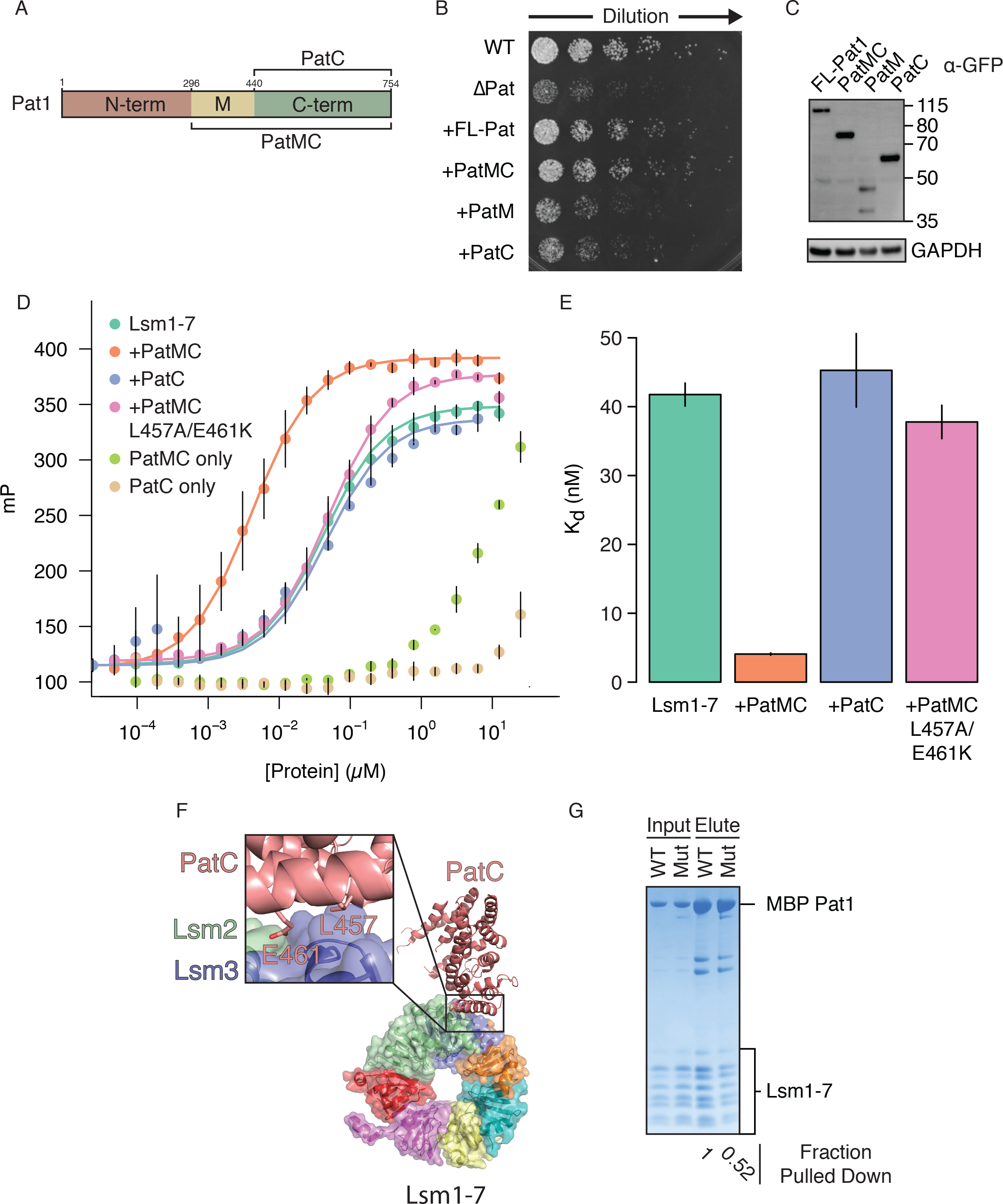
Pat1 makes a bipartite interaction with Lsm1-7 to enhance RNA binding. **A**, Schematic of Pat1 domains. **B**, Plate growth assay of S.*pombe* at 30 °C for ~1.5days with 4-fold dilutions from OD_600_ =0.115 on YES media. FL-Pat1 refers to full length Pat1. All proteins were expressed as C-terminal superfolderGFP fusions under control of the endogenous promoter and with the Ura4 3’ UTR **C**, Western blot for expression of different Pat1 constructs detected with a GPF antibody and GAPDH as a loading control. Molecular weights in kDa are shown on right. **D**, Lsm1-7 binding 5’ FAM-rA15 RNA (Lsm1-7, n=6; +PatMC, n=4, +PatC and +L457A/E461K, n=2) monitored by fluorescence polarization. All Pat1 constructs are N-terminal His_6_MBP fusion proteins. **E**, Equilibrium dissociation constants determined from fits in **D** with standard deviation. **F**, Crystal structure of PatC interaction with Lsm1-7 (PDB 4C8Q) with magnification of PatC region contacting Lsm2 and Lsm3. Residues are numbered according to fission yeast and are conserved. **G**, MBP pull-down assay using 10 *µ*M Lsm1-7 with stoichiometric amounts of MBP-Pat1 construct as input. Mut refers to the *Sp*PatMC L457A/E461K variant.

To better understand how individual domains of Pat1 function in complex with Lsm1-7, we purified Maltose Binding Protein (MBP) fusions of different fission yeast Pat1 constructs and queried if they could enhance RNA binding of the Lsm1-7 complex. We used fluorescence polarization with an rA15 probe (termed oligoA), which mimics a deadenylated tail, to measure RNA binding and examined which regions of Pat1 could enhance the association of Lsm1-7 with RNA. Addition of MBP-PatMC enhanced the RNA binding of Lsm1-7, while PatC showed no effect (Fig. 1D,E). Importantly, PatMC alone bound oligoA RNA weakly, indicating the enhancement of RNA binding was due to enhancement of Lsm1-7 RNA binding. This effect was specific to oligoA, as Pat1 did not enhance binding to a U15 probe (Fig. S1A-C). To test if PatC’s failure to enhance RNA binding of Lsm1-7 was simply due to an inability to form a PatC/Lsm1-7 complex, we incubated PatC and Lsm1-7 together and were unable to observe a stable complex by analytical size exclusion chromatography (Fig. S2A,B). This indicates that the PatC/Lsm1-7 interaction is likely weak and that the middle domain of Pat1 is required to enhance RNA binding of Lsm1-7 in part by promoting formation a stable heterooctameric complex.

Previous crystal structures have identified conserved residues on PatC that contact the Lsm2/3 subunits, and when mutated, lead to loss of complex formation and defects in mRNA degradation in budding yeast (Fig. 1F) (27, 28). Introducing these mutations into *Sp*PatMC (L457A/E461K) minimally affected its association with Lsm1-7, as shown by pull-down analysis, but completely abrogated its ability to enhance Lsm1-7 RNA binding (Figs. 1D,E,G). This suggests that PatC must make specific interactions with Lsm2/3 to promote RNA binding and that the middle domain of Pat1 is required to form a stable complex with Lsm1-7. Furthermore, the inability of PatMC L457A/E461K to promote Lsm1-7 RNA binding suggests the middle domain is necessary, but not sufficient, to enhance function of the complex. We conclude that interactions of the middle and C-terminal domains of Pat1 with the Lsm1-7 ring are required to enhance RNA binding.

### Short-linear motifs in the middle domain of Pat1 bind Lsm1-7 and enhance RNA binding

The middle domain of Pat1 could enhance RNA binding of Lsm1-7 by directly binding RNA, promoting a stable complex formation with Lsm1-7, or through both mechanisms. We first sought to identify which domain of Pat1 could associate with Lsm1-7. A construct containing most of the middle domain of Pat1, specifically residues 296-431, was necessary and sufficient to interact with Lsm1-7, while PatC was unable to form a stable complex by pull-down analysis (Fig. 2). This suggests the middle domain of Pat1 is responsible for promoting a stable complex with Lsm1-7.

**Figure 2:**
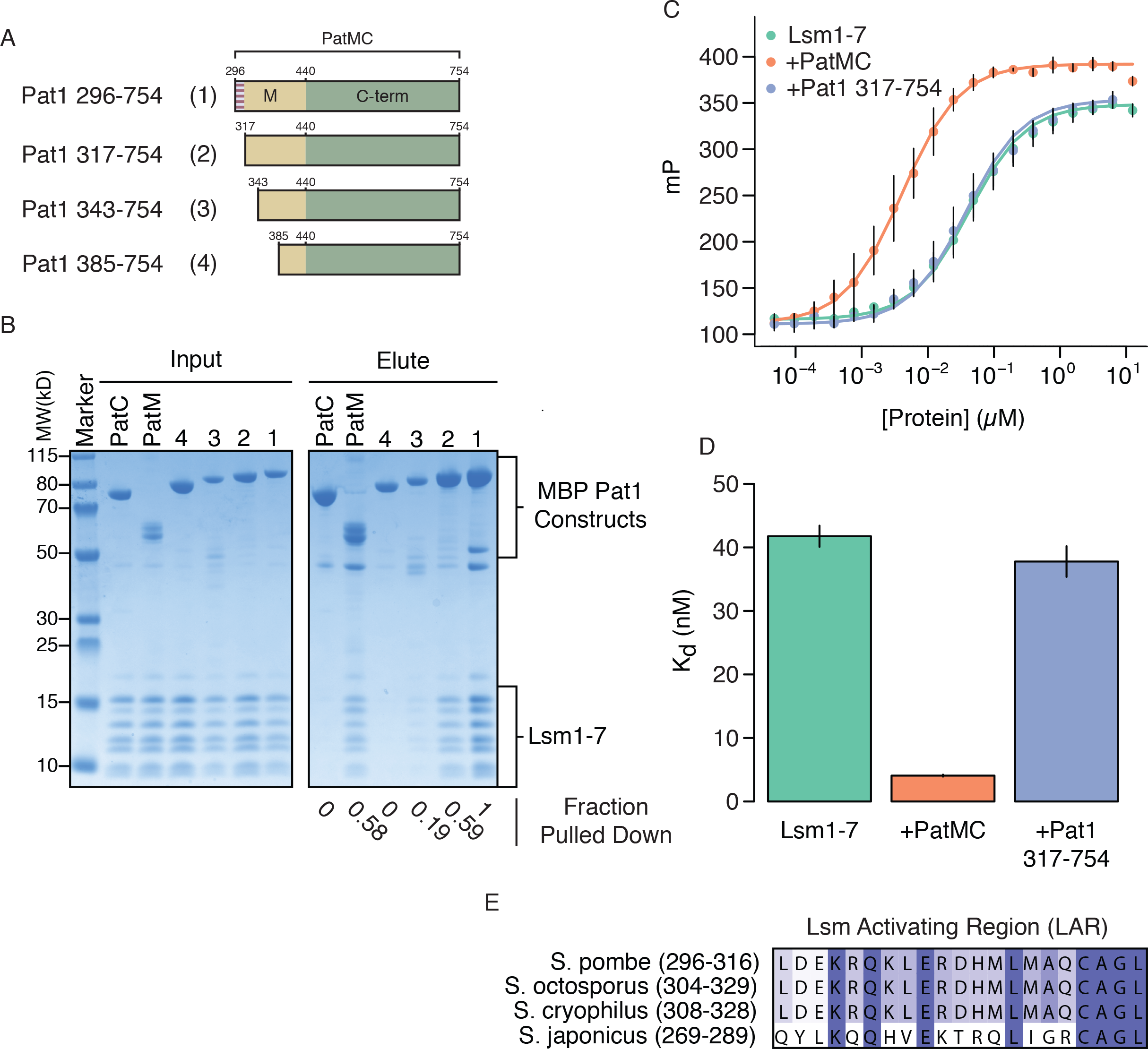
Separate regions of the middle domain of Pat1 interact with and enhance RNA binding by Lsm1-7. **A**, Schematic of different MBP-Pat1 constructs used in pulldown and binding assays. The Lsm Activation Region (LAR) is shown in stripes. **B**, Pulldown of 10*µ*M Lsm1-7 with stoichiometric amounts of specified MBP-Pat1 construct. Numbers refer to schematic in **A**. **C**, Fluorescence polarization of 5’ FAM-rA15 binding to reconstituted MBP-Pat1:Lsm1-7 complexes with standard deviation (Lsm1-7, n = 6; +MBP-PatMC; n=4, +MBP Pat1 317-754; n=4). All conditions contain stoichiometric amounts of Pat1/Lsm1-7. **D**, Dissociation constants of determined from fit to **C** with error bars representing standard deviation. **E**, Sequence alignment of the LAR in different *Schizosaccharomyces* species.

We wanted to distinguish if the middle domain was only involved in protein-protein interactions or had additional roles in promoting RNA binding. To test these possibilities, we purified a series of N-terminal truncations of Pat1 and surveyed their ability to associate with Lsm1-7 by pull-down assays (Fig. 2A,B, truncations 1-4). Deletion of residues 296-431 abolished the Pat1/Lsm1-7 interaction, indicating that these residues are involved in associating with Lsm1-7 (truncation 3). This identifies residues in the middle domain of Pat1 that are required to form a stable complex with Lsm1-7.

Because we identified two N-terminal truncations of Pat1 that retained the ability to interact with Lsm1-7 to similar extent, we tested if these could enhance RNA binding of the PatMC/Lsm1-7 complex (Fig. 2A,B, truncations 1 and 2). If the sole function of the middle domain is to promote association with Lsm1-7, then all N-terminal truncations that interact with Lsm1-7 should enhance RNA binding. Contrary to this expectation, we found Pat1 317-754 was unable to stimulate RNA binding (Fig. 2C,D). This suggests that PatMC contains a region that is dispensable for PatMC/Lsm1-7 complex formation but essential for enhancing RNA binding. The binding isotherm for Pat1 317-754 is identical to the Lsm1-7/PatMC L457A/E461K mutant complex, which can also interact with Lsm1-7 but fails to enhance RNA binding (Fig. 1B,C,E). We therefore reason that residues 296-316 of Pat1 must be involved in stabilizing a productive complex that can bind RNA with high affinity, which we term the ‘Lsm Activation Region’ (LAR). The LAR is highly conserved in the *Schizosaccharomyces* genus, which is characteristic of short linear motifs that can be rapidly transferred between proteins during evolution (Fig. 2E) (18). Metazoans have gained additional residues in this region, though we envision the LAR is still required for Lsm1-7 activation in higher order eukaryotes (Fig. S3).

Our pulldown analysis and functional studies suggest that residues 317-342 in the middle domain of Pat1 are important for promoting a stable PatMC/Lsm1-7 complex. Sequence alignments reveal this region is conserved in Pat1 from yeast to humans (Fig. 3A). To determine if this motif is necessary for the interaction of Pat1 with Lsm1-7, we constructed series of internal PatMC deletions by replacing intervening residues with a flexible (GS)_3_ linker (Fig. 3B). Because appending the conserved motif containing amino acids 317-342 of the middle domain to PatC stabilized the interaction with Lsm1-7 (Fig. 3C), we refer to this region of Pat1 as the Lsm binding motif (LBM). In contrast, deletion of the LAR was dispensable for Pat1/Lsm1-7 interactions. While we cannot exclude that additional regions of Pat1 help stabilize the Pat1/Lsm1-7 complex, this data suggests the LBM helps mediate the protein-protein interaction with the Lsm1-7 ring. We conclude the middle domain of Pat1 contains two short linear motifs that may function separately in stabilizing the PatMC/Lsm1-7 complex and a conformation that is productive for RNA binding.

**Figure 3:**
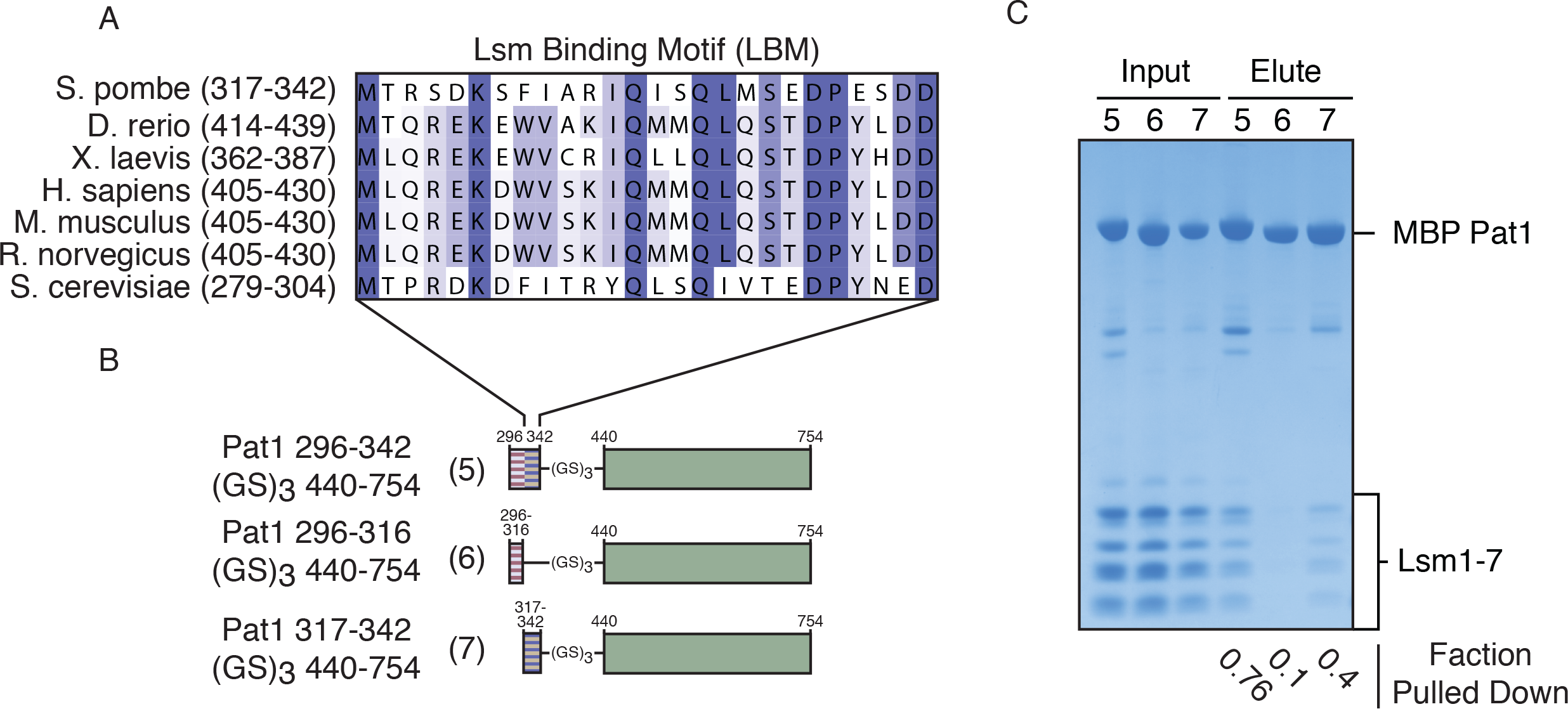
A short linear motif in the middle domain of Pat1 mediates protein-protein interaction with Lsm1-7. **A**, Conservation of the LBM in the Pat1 middle domain between different species. **B**, Schematic of constructs used. Residues were deleted and replaced with a flexible six amino acid (GS)_3_ linker. The Lsm Activation Region (LAR) and Lsm Binding Motif (LBM) is shown in red or purple stripes, respectively. **C**, Pulldown of 10*µ*M Lsm1-7 with stoichiometric amounts of MBP-PatMC internal deletions. Numbers refer to schematic in **A**.

### The middle and C-terminal domains of Pat1 activate Dcp1/Dcp2

Numerous genetic experiments have demonstrated that the middle and C-terminal domains of Pat1 are required to enhance decapping *in vivo*, but the molecular mechanism has remained unclear (28, 36). Dcp2 contains a catalytic NUDIX hydrolase domain that cleaves the m7G cap, as well as a disordered C-terminal extension that is replete with regulatory elements such as inhibitory motifs and protein-protein interaction motifs. While we could not purify full length Dcp2, a C-terminally extended Dcp2 containing up to 261 additional amino acids recapitulates the regulatory elements in the full-length C-terminal extension (Fig. 4A) (16, 17). The C-terminal domain of budding yeast Pat1 was crystallized with a helical leucine rich motif (HLM) of Dcp2, suggesting a mechanism by which Pat1 could promote decapping (41). Accordingly, we tested the ability of Pat1 to activate the decapping holoenzyme, Dcp1/Dcp2 *in vitro* using a C-terminally extended Dcp2 that contains two HLMs (Dcp1/Dcp2 1-318, termed Dcp1/Dcp2_HLM1/2_) (Fig. 4A) (41, 42). We chose this Dcp1/Dcp2 construct for initial studies to examine how Pat1 binding Dcp2 could function independent of inhibitory elements in the C-terminal extension of Dcp2. PatMC was able to enhance the activity of Dcp1/Dcp2_HLM1/2_, while PatC had little effect on activity (Fig. 4B). This suggests that both the middle and C-terminal domain of Pat1 cooperate to directly activate decapping.

**Figure 4:**
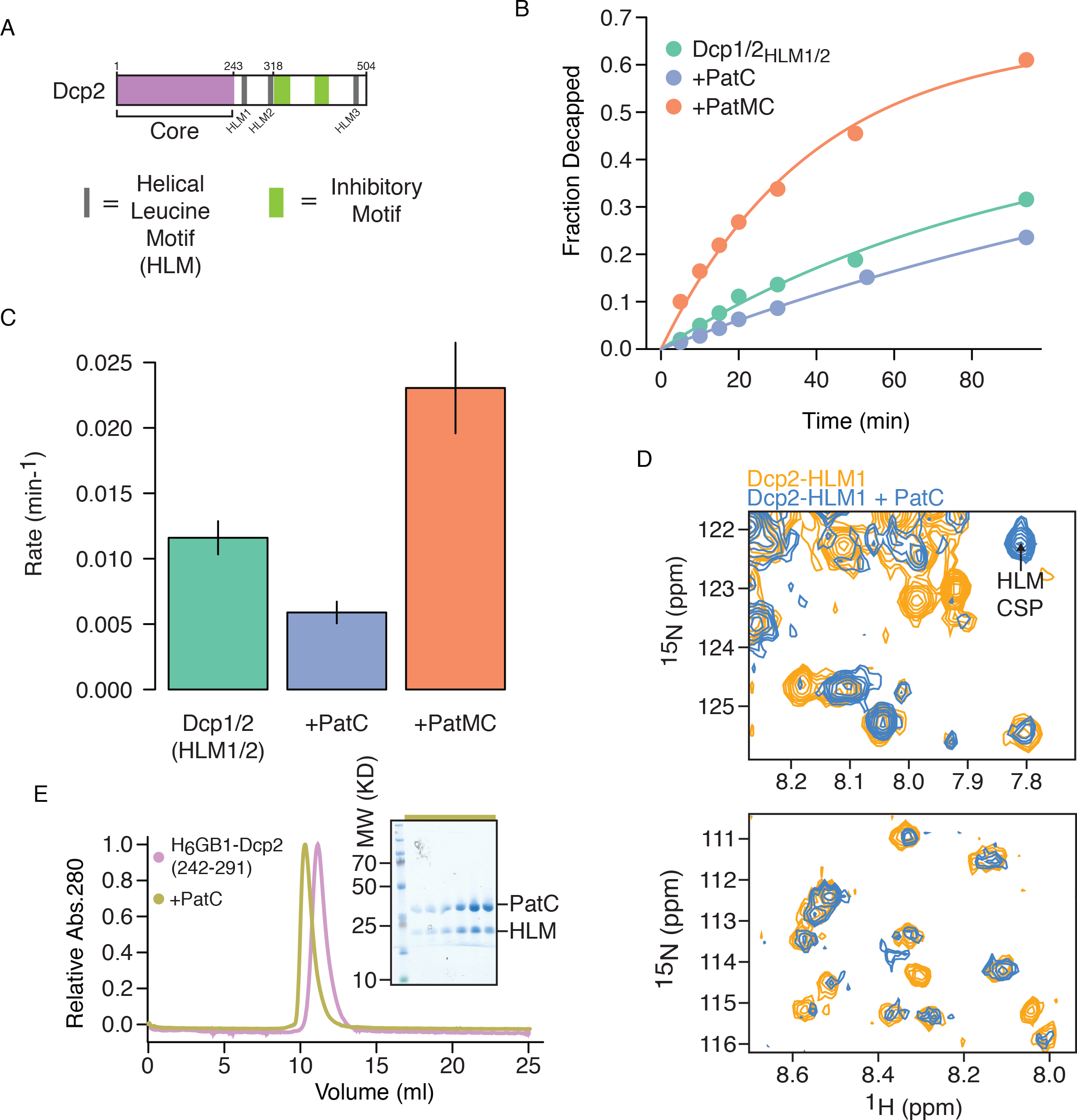
Pat1 directly binds and activates C-terminally extended Dcp2 constructs. **A**, Schematic of domains of Dcp2. The structured core that hydrolyzes cap is shown in purple. Gray boxes represent helical leucine motifs and green boxes represent inhibitory elements. **B**, Representative decapping of an RNA with 150nM Dcp1/Dcp2_HLM1/2_ and 10*µ*M of MBP-tagged Pat1 construct. **C**, Rates determined from **B** displayed with standard deviation (n=2). **D**, Magnified views of regions of interest in ^15^N-^1^H HSQC of 100*µ*M Dcp2_HLM1_ with or without 150*µ*M PatC. Full spectrum is in Figure S1. **E**, Analytical size exclusion chromatography of 35*µ*M H_6_-GB1-Dcp2 242-291 alone (pink) or with a stoichiometric amount PatC (gold). Insert is gel corresponding to peak in gold trace. The helical leucine motif sequence is residues 257-264 of Dcp2.

Although PatC was unable to stimulate decapping on its own, we next asked if it could directly associate with HLMs of Dcp2 using ^15^N HSQC NMR spectroscopy. We purified ^15^N labelled Dcp2 containing a single HLM (Dcp2 1-266, termed Dcp2-HLM1, Fig. 4A), and determined if any chemical shift perturbations (CSPs) occurred in the presence of PatC. Upon addition of PatC, we observed losses and reappearances of resonances in Dcp2-HLM1 but not in the Dcp2 catalytic core alone (Dcp2 1-243, termed Dcp2_Core_) (Figs. 4D & S4). This indicates that PatC directly binds HLMs in Dcp2. Furthermore, HLM-1 of Dcp2 was sufficient for binding PatC, assayed by NMR CSP analysis and copurification by size-exclusion chromatography (Figs. 4E & S5A-D). We conclude that fission yeast PatC is sufficient to interact with Dcp2 by binding HLMs, but both the middle and C-terminal domains are required to activate decapping (41).

To gain insight into how Pat1 activates decapping by Dcp1/Dcp2 in the absence of inhibitory elements, we determined the kinetic parameters of Dcp1/Dcp2_HLM1/2_ either alone or in the presence of MBP-PatMC using a single turnover decapping assay (Fig. 5A). We used saturating PatMC in all experiments to ensure complete formation of a PatMC/Dcp1/Dcp2 complex at all measured concentrations. While we observed minimal effects on k_max_, we saw a ~3-fold reduction in K_m_ (Fig. 5E-F). This suggests that PatMC can bind to HLMs in Dcp2 to enhance substrate binding of the decapping complex.

**Figure 5:**
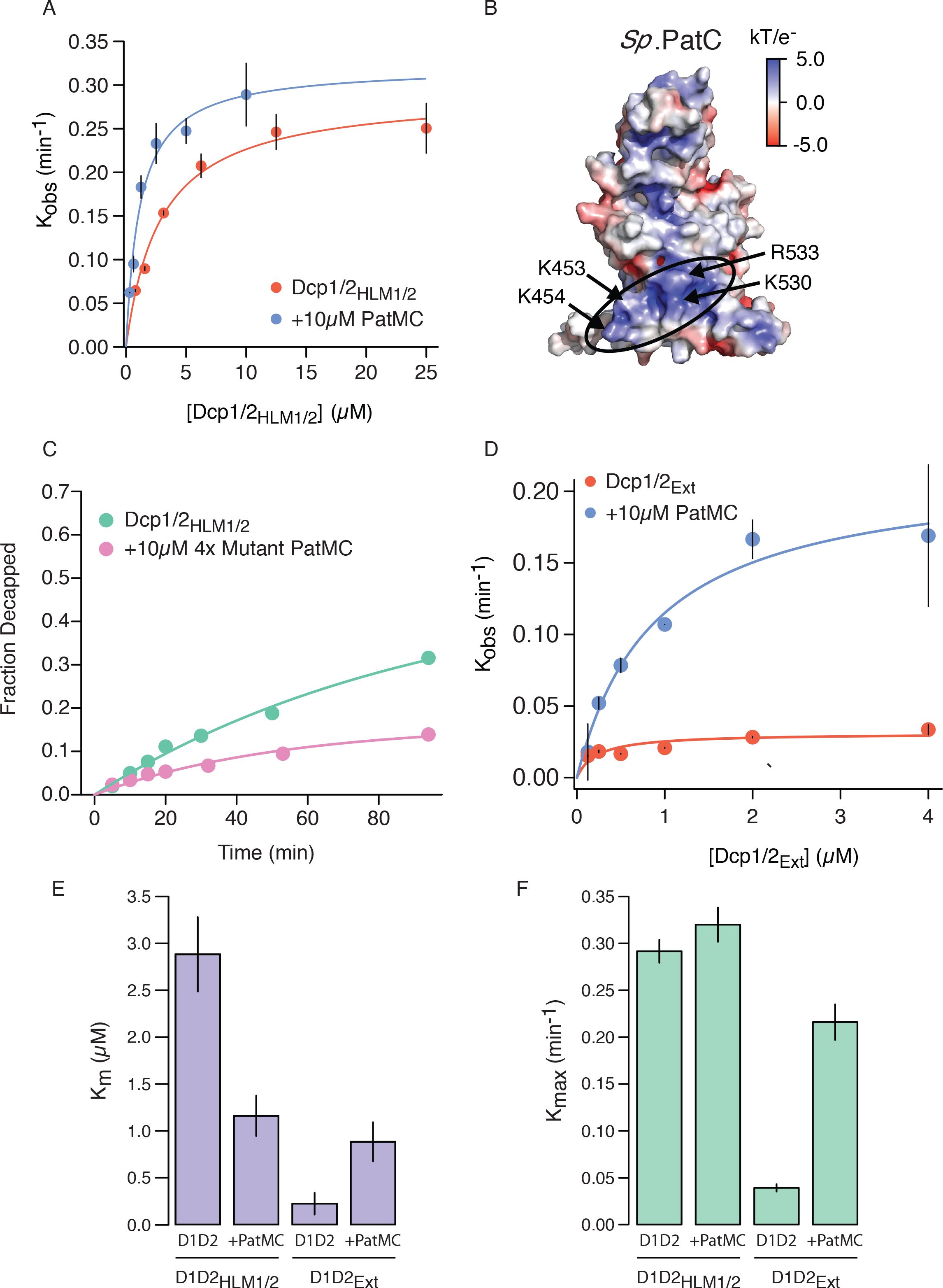
The middle domain differentially activates C-terminally extended Dcp2 constructs. **A**, Single-turnover kinetic analyses of Dcp1/Dcp2_HLM1/2_ alone or in the presence of 10*µ*M MBP-PatMC (n=2)**. B**, Electrostatic surface representation of model of *Sp*PatC based on PDB 5LMG using modeler (47) and APBS in pymol (Schrödinger). Basic patch is circled, while residues mutated are indicated with arrows. **C**, Representative decapping assay of an RNA with 150nM Dcp1/Dcp2_HLM1/2_ and 10*µ*M 4× mutant MBP-PatMC (K453A/K454A/K530A/R533A). **D**, Single-turnover kinetic analyses of autoinhibited Dcp1/Dcp2_Ext_ alone or in the presence of 10*µ*M MBP-PatMC (n=2). **E-F**, K_M_ and k_max_ values calculated from **A** and **D** for Dcp1/Dcp2_HLM1/2_ or Dcp1/Dcp2_Ext_ alone or in the presence of 10*µ*M MBP-PatMC shown with standard deviation.

While *Sp*PatC had no effect on decapping *in vitro*, we hypothesized that, like the Lsm1-7 complex, both the middle domain and C-terminal domains of Pat1 may cooperate to enhance the activity of Dcp1/Dcp2. Studies of human PatC identified a basic patch that is required for mRNA decapping and decay (38). Modeling of *Sp*PatC onto existing crystal structures reveals a similar basic patch to that identified in humans, which we reasoned may be involved in activating Dcp2 (Fig. 5B). Mutation of these four basic residues to alanine (K453A/K454A/K530A/R533A, referred to as 4× Mutant) in *Sp*PatMC abolished its ability to activate Dcp1/Dcp2 (Fig. 5C). Because the middle domain and residues on the C-terminal domain of Pat1 are both required to activate decapping, these domains may form a bipartite RNA-binding surface that can recruit Dcp1/Dcp2 to substrate. This suggests that PatC can bind HLMs and cooperate with the middle domain of Pat1 to increase the affinity of Dcp1/Dcp2 for substrate RNA.

In addition to HLMs, the C-terminal region of Dcp2 contains inhibitory elements that promote nonproductive binding of Dcp1/Dcp2 to substrate to limit catalysis (Fig. 4A). Other HLM binding proteins, such as Edc3, have been shown to relieve autoinhibition (17). To interrogate how PatMC activates an autoinhibited Dcp1/Dcp2, we determined the kinetic parameters of an extended Dcp1/Dcp2 construct that contains both inhibitory elements and HLMs (Dcp1/Dcp2 1-504, termed Dcp1/Dcp2_Ext_), either alone or with saturating amounts of PatMC. Unlike the Dcp1/Dcp2_HLM1/2_, PatMC activated the catalytic step of decapping, while also decreasing the affinity of Dcp1/Dcp2_Ext_ for substrate (Fig. 5A, C-D). This suggests that Dcp1/Dcp2_Ext_ binds nonproductively to RNA, and PatMC activates catalysis at the expense of substrate binding. This is in agreement with HLM binding proteins relieving autoinhibition to promote decapping, suggesting this may be a general model for proteins that bind the C-terminus of Dcp2. We conclude Pat1 uses its middle and C-terminal domains to alleviate autoinhibition and enhance substrate binding of the Dcp1/Dcp2 decapping enzyme complex.

## DISCUSSION

Pat1 works together with Dcp1/Dcp2 and the Lsm1-7 complex to promote bulk 5’-3’ decay on numerous transcripts in yeast (12). Our work reveals how Pat1 interacts with and activates separate steps in this pathway. First Pat1 enhances the RNA binding of Lsm1-7 complex using a bipartite interaction surface comprised of its middle and C-terminal domain. Second, Pat1 promotes decapping by Dcp1/Dcp2 using both domains to affect RNA binding and alleviate autoinhibition. Taken together, these results provide a mechanistic framework for how Pat1 can enhance binding and activity of factors that act on the 5’ and 3’ ends of mRNA to promote bulk decay (Fig. 6).

**Figure 6:**
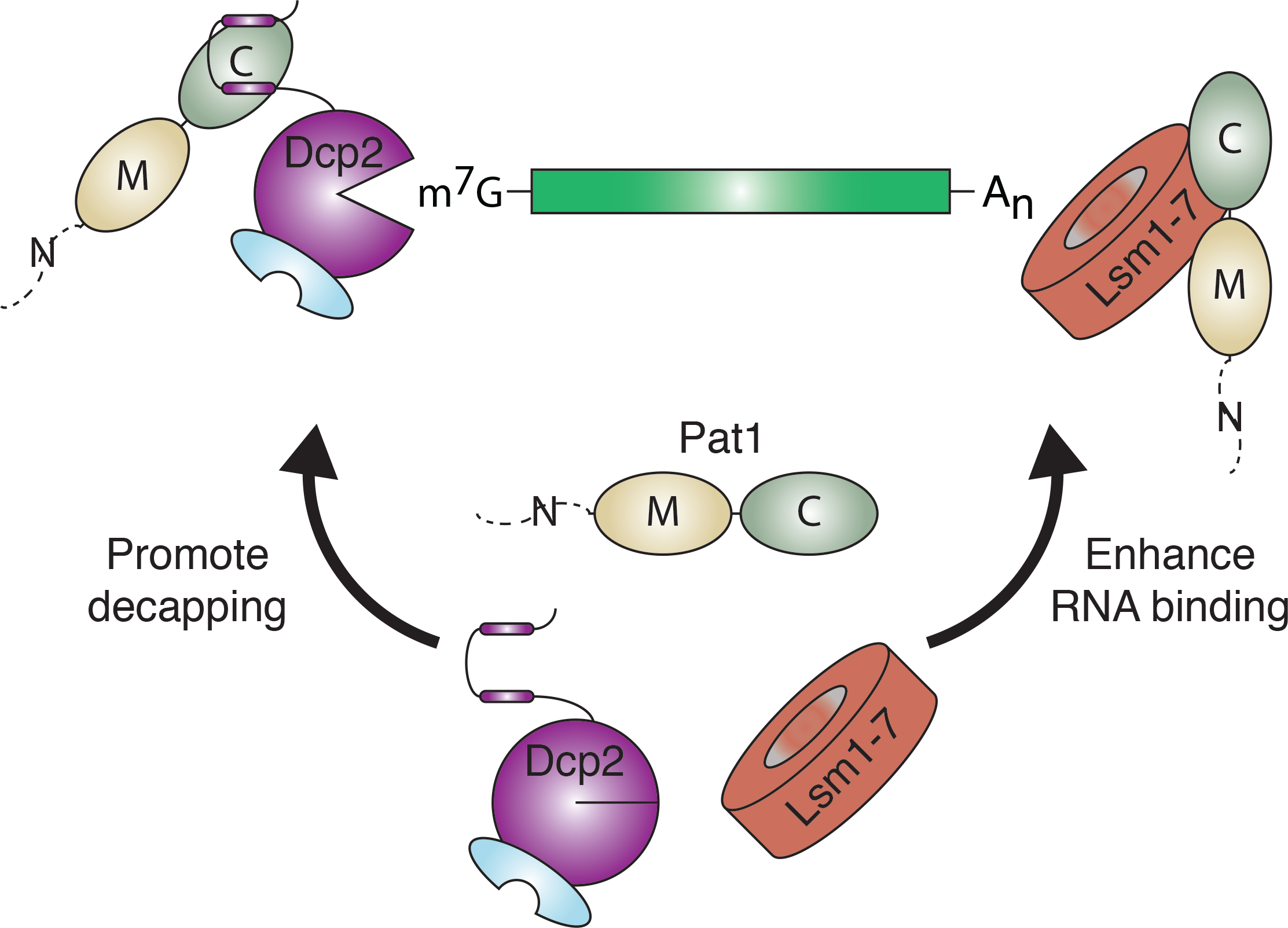
Model for how Pat1 interacts with and activates multiple RNA decay factors to establish 5’ decapping. **A**, Pat1 interacts with and activates decay factors at both 5’ and 3’ end of the mRNA to promote decapping.

Pat1 enhances binding of the Lsm1-7 complex to oligoA RNA 10-fold using bipartite interactions of its middle and C-terminal domains (Fig. 1). This requires short-linear motifs within the middle domain of Pat1, termed the Lsm binding motif (LBM) and Lsm activating region (LAR), that mediate complex formation with Lsm1-7 and enhance oligoA binding (Figs. 2 & 3). Interactions of the C-terminal domain of Pat1 with Lsm2/3, as predicted by crystallographic analyses, are necessary for assembly of a stable complex between Pat1, the Lsm1-7 ring and RNA. Pat1 constructs lacking the Lsm activating region (LAR) still interact with Lsm1-7 but fail to enhance RNA binding. Therefore, protein interactions required for formation of a stable Pat1 and Lsm1-7 complex can be separated from the ability of Pat1 to enhance RNA binding of Lsm1-7. The middle domain has a role beyond the promoting the interaction between Pat1 and Lsm1-7 and is likely involved in stabilizing a conformation of Lsm1-7 that can bind RNA with high affinity. The molecular details of how the middle domain of Pat1 interacts with the Lsm1-7 ring and enhances RNA binding will require high-resolution structural studies.

A mechanistic description of how the Pat1/Lsm1-7 complex is recruited to target mRNAs remains poorly understood. In budding yeast, Pat1/Lsm1 bind the 3’ end of the mRNA and promote degradation of inefficiently translated mRNAs (12, 30). In S.*pombe* and higher order eukaryotes, however, an additional 1-3 uridines are added to a subset of transcripts after deadenylation but prior to decapping, which are enriched when Lsm1 is deleted (19, 20). Understanding how Pat1/Lsm1-7 recognition of mRNAs is coupled to different tail modifications in cells remains a challenge for the future.

Multiple decapping activators bind the C-terminal regulatory region of Dcp2 to activate decapping and ensure transcript specificity in cells. We show that Pat1 directly activates decapping by binding helical leucine rich motifs (HLMs) in the C-terminal regulatory region to enhance Dcp1/Dcp2 RNA binding and alleviate autoinhibition (Figs. 4–5). This mechanism is reminiscent of other HLM binding proteins, such as Edc3, and may be a general mechanism by which HLM binding proteins promote decapping. Importantly the middle domain and structured C-terminus are both required for these effects. This is consistent with early work showing the middle domain is required to activate decapping in yeast and recent structural work showing C-terminal domain binds Dcp2 (16, 41). Taken with our biochemical characterization of the Pat1/Lsm1-7 interaction, these observations suggest that both the middle and C-terminal domains of Pat1 are required to activate distinct steps of mRNA decay. It is likely that deletion of the middle domain has a strong defect in decapping because of reduced Lsm1-7 RNA binding and activity of Dcp1/Dcp2 on substrate mRNA.

Our work demonstrates how conserved short-linear motifs of the unstructured middle domain of *Sp*Pat1 enhance Lsm1-7, which may be applicable to nuclear Lsm2-8 complexes. Recent genome-wide analysis of the vertebrate orthologue of Pat1 (Pat1b/PatL1) suggest it functions with the Lsm2-8 complex during pre-mRNA splicing in addition to its established role with Lsm1-7 in mRNA decay. Ablation of Pat1b promotes skipping of exons with weak donor site sequences, in addition to increasing the steady-state levels of AU rich containing mRNAs. Knockdown of Pat1b reduces formation of the U6 snRNP containing Lsm2-8 and SART3 (an orthologue of yeast Prp24). Furthermore, Pat1b is localized in nuclear Cajal bodies as well as cytoplasmic processing bodies (43). The unstructured middle domain of Pat1 may endow it with structural plasticity, allowing it to function as a versatile chaperone for RNP assembly during bulk 5’-3’ mRNA decay and pre-mRNA splicing.

## ACKNOWLEDGEMENTS

We thank RA Greenstein for help with construction of yeast strains, members of the Gross lab for discussion and Jeffrey S. Mugridge and Krister J. Barkovich for comments on manuscript. R.W.T is supported by a Genentech Foundation predoctoral fellowship and UCSF discovery fellowship. This work was supported by US National Institutes of Health grant R01 GM078360 to J.D.G.

## AUTHOR CONTRIBUTIONS

J.H.L and J.D.G conceived of study. J.H.L performed experiments. J.H.L and R.W.T performed NMR. J.H.L and J.D.G wrote and edited manuscript.

## MATERIALS AND METHODS

### RNA transcription and capping

A 340 nucleotide RNA containing a short oligo A tail of 15 nucleotides and corresponding to the MFA2 gene of budding yeast was *in vitro* transcribed and capped according to previous protocols (44). Briefly, RNA was transcribed using T7 RNA polymerase and purified on a 6% denaturing polyacrylamide gel. RNA was excised from gel and extracted before being further purified by phenol:chloroform and ethanol precipitation. RNA was capped with α-^32^P GTP using vaccinia virus capping enzyme and separated by gel purification, phenol:chloroform extraction and ethanol precipitation as previously described (44).

### Decapping Kinetics

Decapping assays with Dcp1/Dcp2 and cofactors were performed at room temperature in a buffer containing 150mM NaCl, 20mM HEPES pH = 7, 1mM DTT, 5mM MgCl_2_ with 0.3μg/μl Acetylated Bovine Serum Albumin. Reactions were performed under single-turnover conditions, where enzyme (Dcp1/Dcp2) is in excess over substrate. The concentration of MBP-Pat1 constructs was 10 *µ*M MBP-Pat1 in all experiments. Proteins were mixed at desired concentrations and incubated at room temperature for 30 minutes before initiating the reaction with ^32^P capped-RNA. Time points were taken and quenched in 0.5 M EDTA before being analyzed by Thin Layer Chromatography. Data were phosphoimaged with a Typhoon scanner (GE) and analyzed using ImageQuant software (GE). Rates were obtained by fitting to a first-order exponential or, when applicable, a linear model. Kinetic constants were obtained by increasing Dcp1/Dcp2 concentrations and time-courses as previously described (44).

### Fluorescence Polarization

Fluorescence polarization was performed in 384-well low volume plates (Greiner) in the same buffer used for decapping kinetics. Oligo RNAs with 5’ FAM were ordered from IDT and used at a final concentration of 500 pM. To assemble Pat1/Lsm1-7 complexes, stoichiometric amounts of the Pat1 construct and Lsm1-7 were mixed together on ice for 10 minutes prior to setting up reaction. Reactions were incubated for five minutes before measuring polarization on an Analyst AD plate reader (LJL Biosystems). Equilibrium dissociation constants (K_d_) were fit to the equation for oligo A15:

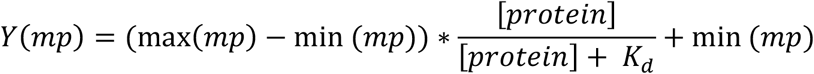

or for U15, when probe concentration was close to K_d_

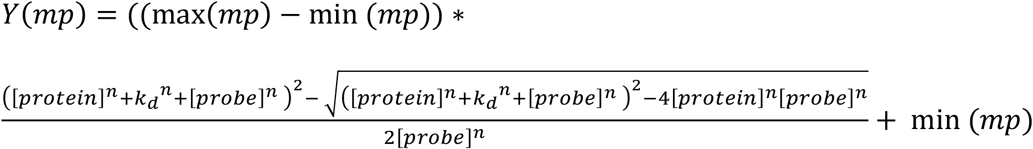

where n represents the Hill coefficient. Plots and fits were generated using in house scripts written in R, which are available upon request.

### Cloning and Protein Purification

All seven Lsm genes were codon optimized for expression in *E.coli* and synthesized as polycistronic gene (IDT). The genes were in order from Lsm1 to Lsm7, where each gene was separated by a 28nucleotide spacer followed by a ribosome binding site. The polycistron was cloned into a vector with an His_6_-TEV site on the N-terminus of Lsm1. Unless stated otherwise, proteins were purified in BL21(DE3)* in LB media. Cells were grown to OD_600_=0.6 at 37 °C. IPTG was added to 1 mM and cells were grown shaking overnight at 17 °C. Cell pellets were harvested by centrifugation and lysed in appropriate buffer. For Lsm1-7, cells were lysed in Buffer A (2 M NaCl, 20 mM HEPES, 20 mM Imidazole, 5 mM βME, protease inhibitor, lysozyme, pH=7.5) by sonication. Lysate was subsequently clarified by centrifugation and the supernatant was bound to Ni-NTA resin at 4 °C for 1 hour. The resin was then loaded into a gravity column and washed with 20 column volumes of Buffer A. All proteins were eluted in Buffer E (250 mM NaCl, 250 mM Imidazole, 20 mM HEPES, 10 mM βME) and tags were cleaved with TEV protease overnight at 4 °C before being loaded onto a HiTrap Heparin column (GE). The heparin column was run at 2 ml/min from a 0.25-1 M NaCl gradient over 20 column volumes. Fractions containing protein complex were collected and further purified by gel filtration using a Superdex 200 16/60 column (GE) equilibrated in SEC buffer (20 mM HEPES pH =7, 150 mM NaCl, 1 mM DTT). Fractions containing protein were concentrated before being flash frozen and stored at −80 °C.

For expression of GB1-HLM1 (residues 242vto 291 of spDcp2) and MBP-Pat1 constructs, cells were lysed in Buffer B (500 mM NaCl, 20 mM Imidazole, 5 mM βME, 20 mM HEPES pH = 7) and bound to nickel resin before being washed with 20 column volumes of Buffer B. Protein was then eluted in with Buffer E, and if necessary, tags were TEV cleaved overnight. We found the MBP tag to be essential for purification of all Pat1 constructs that contained the middle domain, so it was not removed. Elutants were concentrated and further purified by S200 16/60 in SEC buffer. Peak fractions were pooled and concentrated before being flash frozen.

Dcp1/Dcp2_HLM1/2_ or Dcp1/Dcp2_Ext_ was coexpressed and purified by Ni-NTA chromatography as described for MBP-Pat1 except with an additional purification step. After elution from Ni-NTA resin, TEV was added and the protein was cleaved overnight before being loaded onto Streptactin resin (IBA). The column was washed with 20 column volumes of Buffer B and eluted in Buffer B supplemented with 10 mM desthiobiotin. The elution was the concentrated and purified on an S200 16/60 in SEC buffer. Sodium Chloride was added to 300 mM before being concentrated and flash frozen.

### Copurification by Analytical Size Exclusion Chromatography

Samples were mixed at ~35μM in 500μl volume before being filtered and injected onto a GE Superdex 75 10/300. All samples were run in 20mM HEPES pH =7.5, 1mM DTT with either 400 mM NaCl for Lsm1-7+PatC or 150 mM NaCl for PatC+GB1 Dcp2 242-291. Samples were run at 0.35ml/min and peaks were analyzed by SDS-PAGE (Invitrogen).

### NMR

^15^N labelled GB1 Dcp2 242-291 (HLM1), Dcp2 1-243 (Dcp2_core_), and Dcp2 1-266 (Dcp2-HLM1) were expressed in H_2_O-based minimal media with ^15^NH_4_Cl as the sole nitrogen source. For ^13^C ILV labelling of GB1-Dcp2 242-291 precursors were added (Ile: 50mg/L, Leu/Val: 100mg/L) 40minutes prior to induction. IPTG was added to 1mM and proteins were expressed overnight at 18 °C. Cells were lysed in Buffer B and purified by on Ni-NTA resin. For Dcp2 constructs, the tag was cleaved by addition of TEV. Proteins were then purified by size-exclusion chromatography on a superdex 75 16/60 in NMR buffer (21 mM NaH_2_PO_4_, 28.8 mM Na_2_PO_4_, 200 mM NaCL, 100 mM Na_2_SO_4_, 5mM DTT). All ^1^H-^15^N and ^1^H-^13^C HSQC experiments with PatC and Dcp2 were performed with 100 μM Dcp2 and 150 μM PatC in NMR buffer and recorded at 303 K on a Bruker Avance 800 spectrometer equipped with a cryogenic probe.

### Pull downs

All pulldowns were done in SEC buffer. Briefly, 10*µ*M of MBP-tagged Pat1 construct and 10*µ*M Lsm1-7 were mixed and incubated at room temperature for 15 minutes before being added to pre-equilibrated amylose beads (New England Biolabs) for 5 minutes. The beads were spun down at 4000g for 2 minutes and samples were washed four times with 500 *µ*l SEC buffer before being eluted with 30 *µ*l SEC buffer supplemented with 10 mM Maltose. Eluted samples were resolved on 4-12% Bis-Tris SDS-PAGE gel (Invitrogen) and stained with Instant Blue (Expedeon). All input was loaded diluted 8-fold compared with elution to account for unbound Pat1/Lsm1-7 constructs. Fraction pulldown was calculated from the following equation:

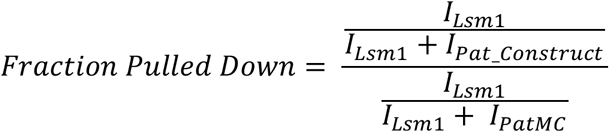

where I_lsm1_ and I_PatMC_ are the intensities of the background subtracted Lsm1 and PatMC intensities, respectively. I_Pat_Construct_ is the intensity of the specified Pat1 construct. This equation normalizes all pulldowns to PatMC/Lsm1-7. All intensities were obtained by quantifying the same size area in ImageJ and background subtracted. Lanes that had Lsm1 intensities below background were adjusted to zero. We chose to quantify only Lsm1 band, as commassie staining is nonlinear and decreases with the molecular weight of the protein.

### Fission yeast strain construction and growth assays

All strains were constructed by knocking in C-terminally tagged superfolderGFP Pat1 constructs into the commercially available pat1::KanMX background (Bioneer). A list of strains used in this study are provided in **Table S1**. All Pat1 constructs were cloned with a C-terminal superfolderGFP tag and Ura4 3’ UTR and cloned into a pUC-19 vector containing 500basepairs upstream and downstream of the Pat1 locus with a Hygromycin resistance cassette (45). Plasmids were linearized with restriction enzymes and knocked into the ∆Pat1 background by standard procedures (46). To verify knock in at the correct genomic locus DNA was obtained by phenol:cholorom extraction and tested for presence of correct genomic insertion by PCR. Colonies containing the knock in at the correct locus were stored as glycerol stocks at −80 °C. For plate growth assays, yeast strains were grown overnight in YES and diluted to OD_600_ = 0.115 and serially diluted 4-fold before being spotted onto YES plates and grown for ~1.5 days at 30 °C.

For western blots, cells were grown overnight and diluted to OD_600_ = 1 the following day. Cells were pelleted by centrifugation and protein was extracted with 2M NaOH/10mM BME supplemented with a protease inhibitor tablet (Roche). 55% Trichloroacetic acid was then added to precipitate protein, and then centrifuged at 16,000rpm for 20 minutes at 4 °C. The supernatant was removed and the pellet was resuspended in HUE buffer (8M Urea, 1%SDS, 50mM EDTA, 10mM BME) and heated to 70 °C for 15 minutes before flash freezing and storing at −80 °C. Western blots for protein expression were performed using antibodies against GFP and GAPDH as a loading control (Proteintech) and detected with StarBright^TM^ 700 goat anti-mouse IgG (Biorad) on a ChemiDoc (Biorad) using the Qdot705 channel.

## SUPPLEMENTAL FIGURE LEGENDS

**Supplemental Figure 1:**
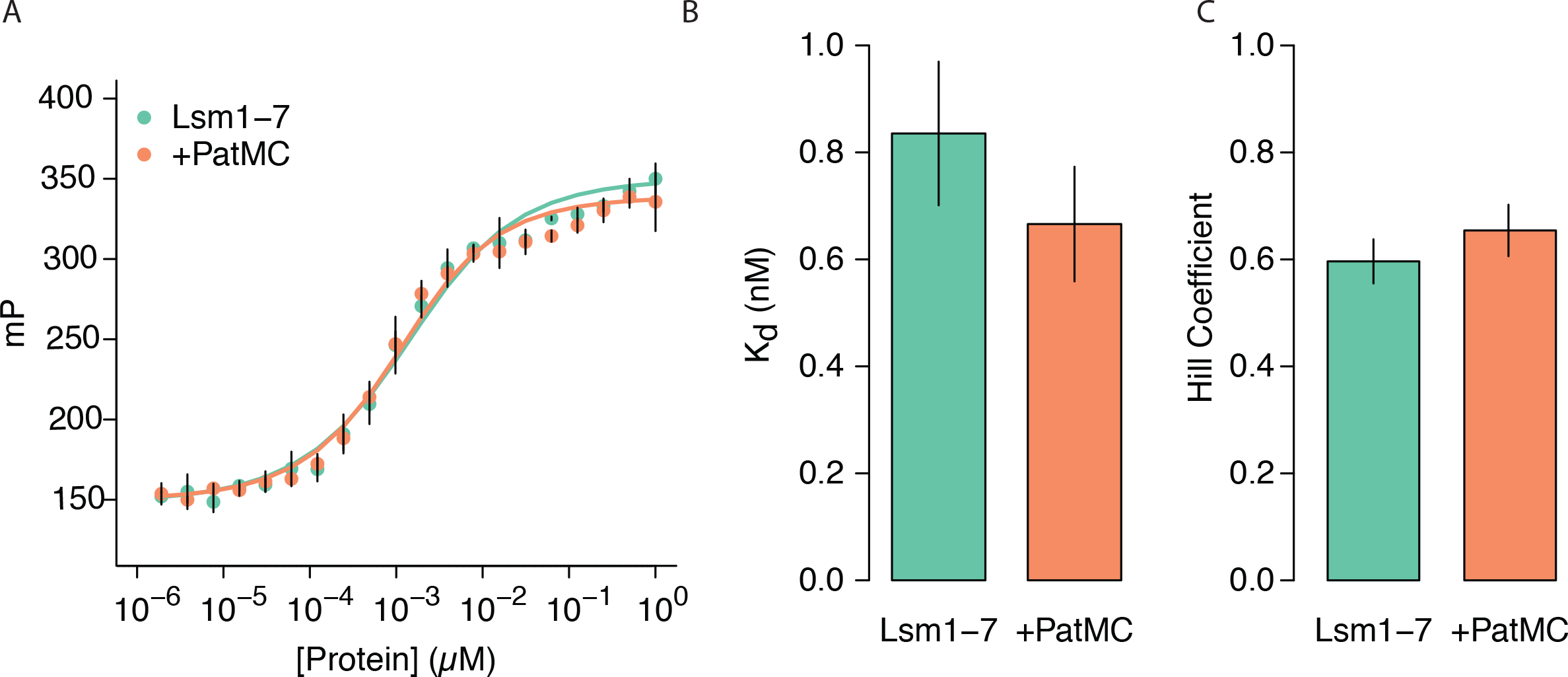
PatMC does not enhance RNA binding for U15: **A**, Lsm1-7 binding 5’ FAM-U15 RNA alone or with H_6_MBP-PatMC (both n=3) monitored by fluorescence polarization. The data was fit to a quadradic binding equation with cooperativity. **B**, Dissociation values from **A** with standard deviations. **C**, Hill coefficients from **A** with standard deviations.

**Supplemental Figure 2:**
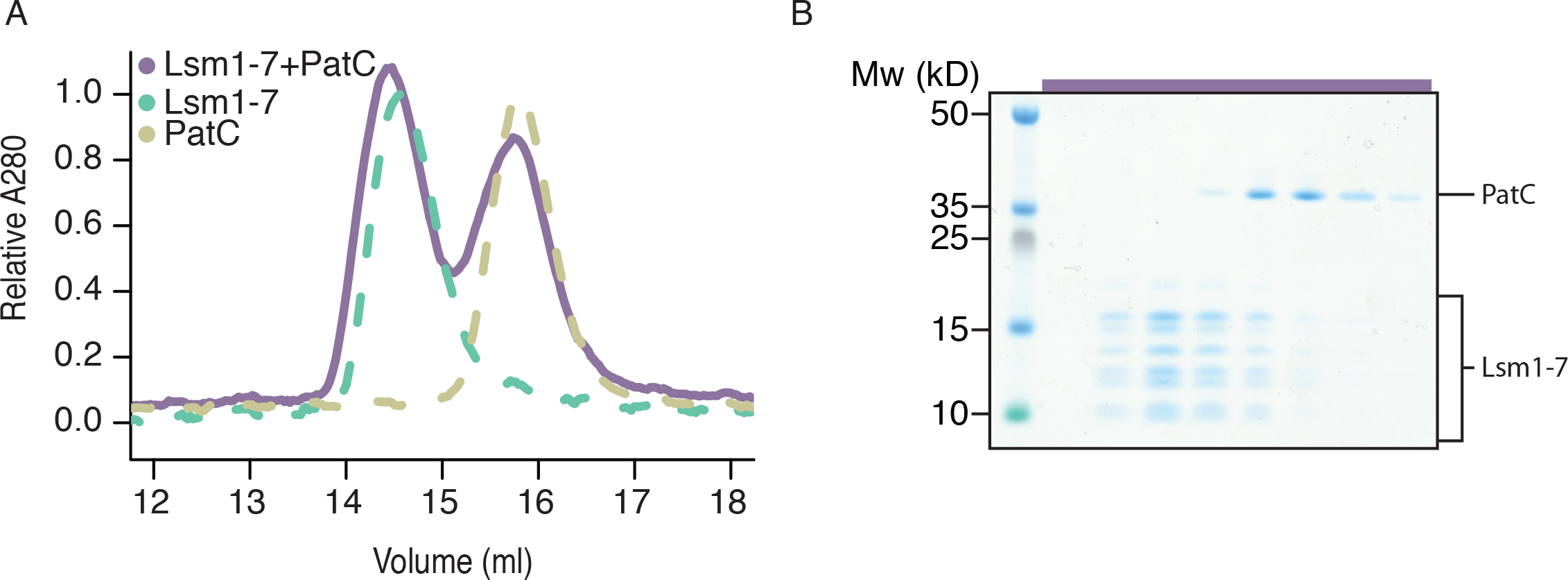
PatC does not stably interact with Lsm1-7. **A**, Analytical size exclusion chromatography of PatC, Lsm1-7, or combined PatC/Lsm1-7. All proteins were at ~25*µ*M and run in a 400mM NaCl buffer. **B**, Gel corresponding to fractions of Lsm1-7 + PatC SEC in Figure 3C. The purple bar on top corresponds to purple chromatogram.

**Supplemental Figure 3:**
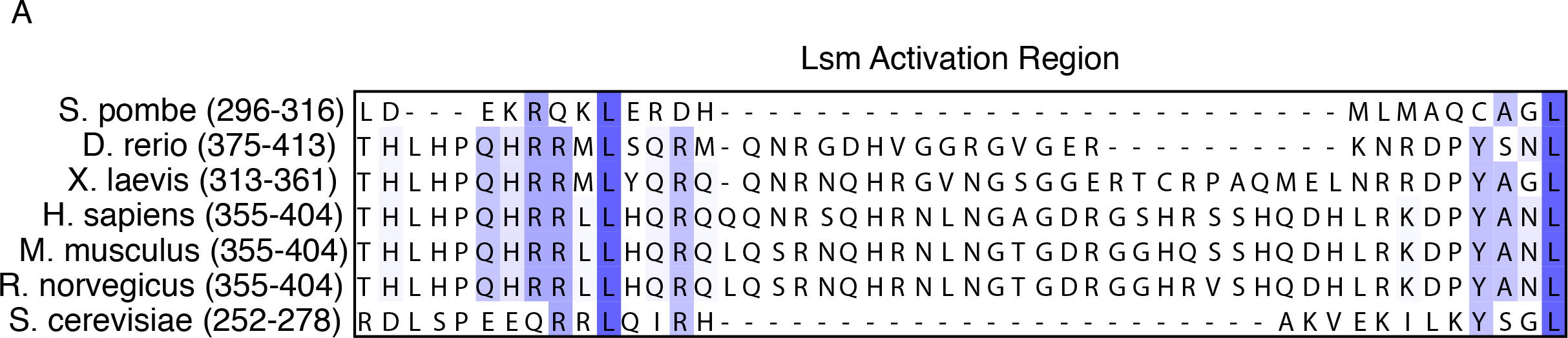
Sequence alignments of the Lsm Activation Region (LAR). **A**, Sequence alignment of different species of the LAR. Metazoans have acquired additional residues that may be functionally important for enhancing the RNA binding of Lsm1-7.

**Supplemental Figure 4:**
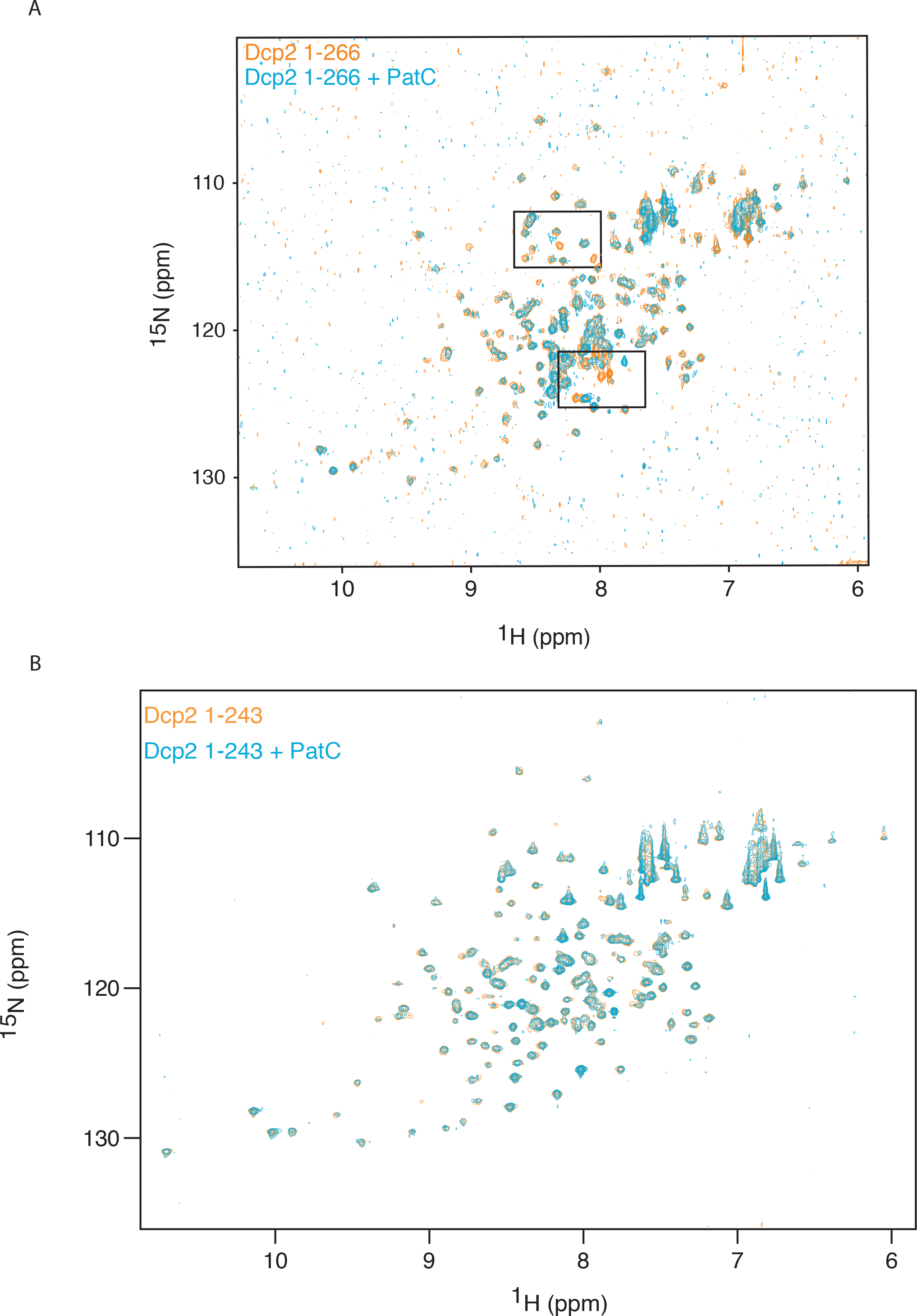
Full ^15^N-^1^H HSQC of Dcp2 constructs ± PatC. **A**, Full spectrum of 100*µ*M ^15^N labeled Dcp2_HLM1_ in the presence or absence of 150*µ*M PatC. Boxes represent magnified views shown in Figure 1D. **B**, ^15^N-^1^H HSQC of 100*µ*M Dcp2_Core_ with or without 150*µ*M PatC.

**Supplemental Figure 5:**
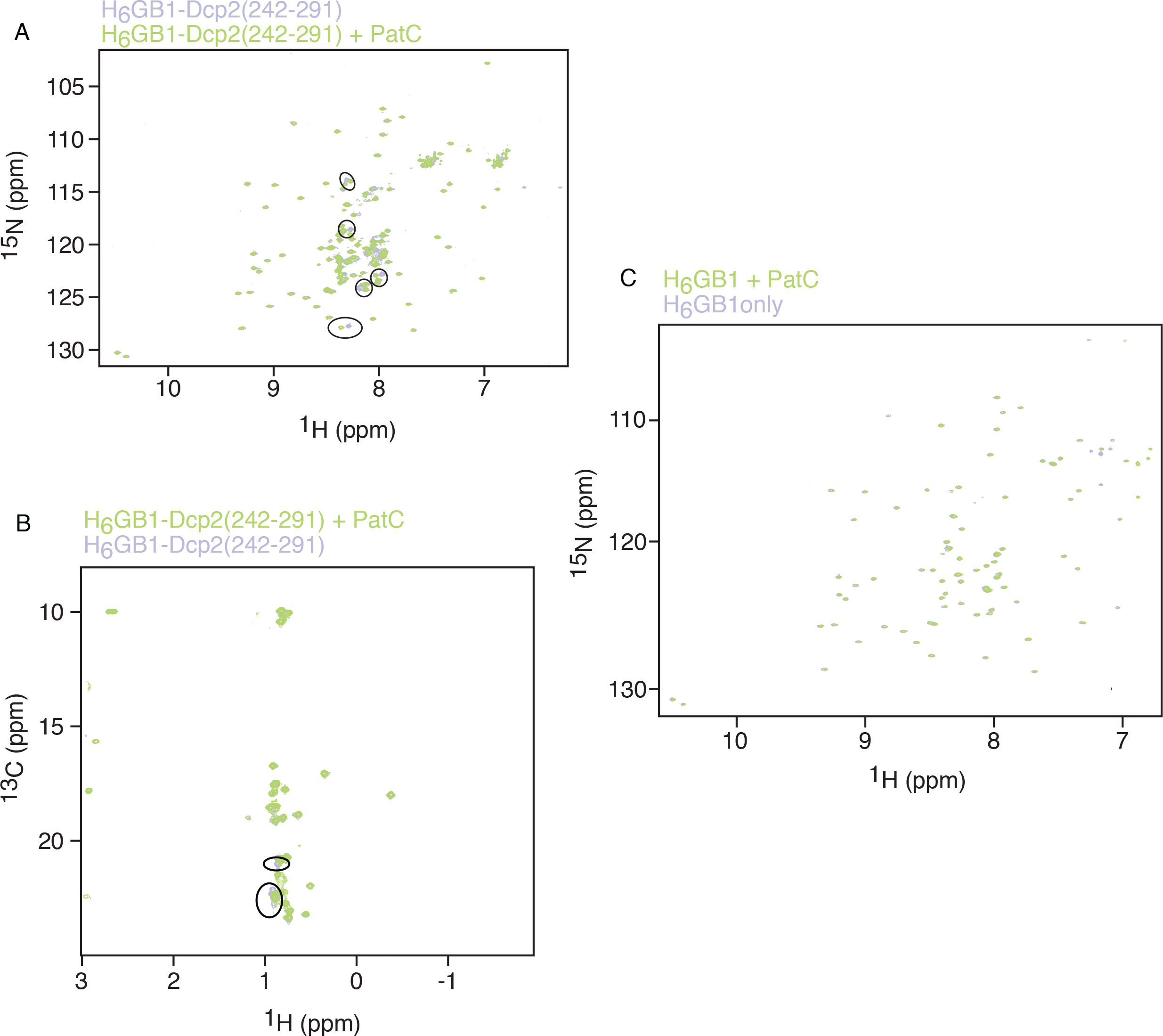
PatC interacts with an isolated HLM. **A**, ^15^N-^1^H HSQC of 200*µ*M uniformly ^15^N labelled H_6_-GB1-Dcp2 242-291 (purple) alone or 100*µ*M H_6_-GB1-Dcp2 242-291 with 200*µ*M PatC (green). Circles indicate sites of chemical shift perturbations. **B**, ^13^C-^1^H HSQC of 200*µ*M ^13^C ILV labeled H_6_-GB1-Dcp2 242-291alone (purple) or 100*µ*M H_6_-GB1-Dcp2 242-291 with 200*µ*M PatC (green). **C**, No chemical shifts of 50*µ*M _15_N-_1_H HSQC H_6_-GB1 in the presence of 100*µ*M PatC (purple). 100*µ*M H_6_-GB1 alone is shown in green.

## SUPPLEMENTAL TABLE LEGENDS

**Supplemental Table 1: S.*pombe* strains used in this study.**

